# How a sticky fluid facilitates prey retention in a carnivorous pitcher plant (*Nepenthes rafflesiana*)

**DOI:** 10.1101/2021.03.13.434712

**Authors:** Victor Kang, Hannah Isermann, Saksham Sharma, D Ian Wilson, Walter Federle

**Author notes:** Corresponding author: V.K. Present address: Department of Bioengineering, Imperial College London, London, UK.

## Abstract

*Nepenthes* pitcher plants live in nutrient-poor soils and produce large pitfall traps to obtain additional nutrients from animal prey. Previous research has shown that the digestive secretion in *N. rafflesiana* is a sticky viscoelastic fluid that is much more effective at retaining insects than water, even after significant dilution. Although the physical properties of the fluid are important for its retentive function, it is unclear how the fluid interacts with insect cuticle and how its sticky nature affects struggling insects. In this study, we investigated the mechanisms behind the efficient prey retention in *N. rafflesiana* pitcher fluid. By measuring the attractive forces exerted on insect body parts moving in and out of test fluids, we show that it costs insects significantly more energy to separate from pitcher fluid than from water. Moreover, both the maximum force and the energy required for retraction increase after the first contact with the pitcher fluid. We found that insects sink more easily into pitcher fluid than water and, accordingly, the surface tension of *N. rafflesiana* pitcher fluid was significantly lower than that of water (60.2 vs. 72.3 mN/m). By analysing the pitcher fluid dewetting behaviour, we demonstrate that it strongly resists dewetting from all surfaces tested, leaving behind residual films and filaments that can facilitate re-wetting. This inhibition of dewetting may be a further consequence of the fluid’s viscoelastic nature and likely represents a key mechanism underlying prey retention in *Nepenthes* pitcher plants.

## 1. Introduction

Pitcher plants are striking examples of plants that have turned carnivorous to supplement the nutrient-poor soils they inhabit. Through their characteristic pitfall traps made from highly modified leaves – a design that has evolved independently at least six times across the kingdom – these plants lure, retain, and finally digest prey [1,2]. The prey, which includes mostly ants but also flying insects [3], are normally capable of climbing up vertical surfaces or flying away from danger, yet they struggle to escape from pitfall traps due to a combination of specialised structural adaptations. In *Nepenthes* pitcher plants (Nepenthaceae), the pitfall traps consist of a lid, a peristome, a slippery zone on the inner wall, and a digestive zone. The lid generally serves to prevent excessive dilution from rainfall, but in some species it also facilitates prey capture [4]. The highly wettable peristome causes insects to slip on a stable water-film and fall into the pitcher [5,6]. Depending on the species, the slippery zone is covered by wax crystals that readily break off to contaminate insect tarsi and also produce fine-scale roughness that impedes insect adhesion and their attempts to climb out [7–9]. The slippery zone can also contain directional microstructures that make it easier for insects to slip into and impede escape from the pitcher [10]. Finally, the walls of the digestive zone are covered in glands for secretion and absorption, but they are unlikely to serve a role in prey retention [11,12].

Although the structural adaptations of pitfall traps help to capture and retain nonflying prey that need to scale the inner wall to escape, they are less suitable against flying insects. Indeed, video recordings of flies falling into containers of water show that they are able to recover and fly away without contacting the sides, which suggests that watery pitcher fluid may be less effective in catching flying insects [13]. Since pitcher plants catch a variety of flying and nonflying insects, it is likely that other mechanisms further enhance their performance [3,14,15]. As insects that fall into the traps land on the digestive fluid, it is possible that the fluid itself can help to maintain the prey. This mechanism would further benefit the plant by prohibiting the escape of nonflying insects as well, which can sometimes overcome the aforementioned structural adaptations and climb out [5]. In several species of *Nepenthes*, including *N. rafflesiana*, *N. hemsleyana,* and *N. gracilis*, significantly more flies and ants were retained in isolated digestive fluid than in water [13,16,17]. Fluid from young *N. rafflesiana* plants were the most effective, retaining 100% of the tested flies and ~90 to 100% of the ants, while less than 20% of the flies and none of the ants were retained in water [16]. Such striking differences in retention rates of pitcher fluid compared to water have also been reported in several members of the Sarraceniaceae family, which independently evolved pitfall traps to catch prey. Experiments using digestive fluid from *Sarracenia flava*, *S. sledgei* (synonym *S. alata*)*, S. drummondii* (synonym *S. leucophylla*), and *Darlingtonia californica* demonstrated that ants sank more rapidly in digestive fluid than in water [18–20]. Importantly, insects rescued at the end of the retention trials survived, which indicate that the high retention rates are unlikely caused by noxious compounds released into the fluid. These findings support the idea of a dual functionality of the ‘digestive’ fluid, where it serves to both retain and digest prey. Thus, to recognise the retentive function of the fluid, we refer to it as pitcher fluid (PF) henceforth.

Despite the evidence for the retentive role of PF, we have yet to fully understand its underlying mechanisms. Researchers have previously focused on two PF properties - viscoelasticity and surface tension - to explain how it may function. Many *Nepenthes* species produce PFs that form long sticky filaments when rapidly extended, which is characteristic of non-Newtonian viscoelastic fluids containing high molecular weight polymers [13,21,22]. In an earlier study, the researchers explored the viscoelastic properties of *N. rafflesiana* PF and suggested that its high apparent extensional viscosity and long relaxation times make it more difficult for a struggling insect to swim in and extract itself from the fluid [13]. Although these findings offer insights into the rheological properties of the fluid, they do not help us understand how the fluid interacts with the insect, and why insects ultimately fail to escape. Furthermore, it is unclear how rheological parameters such as extensional viscosity and relaxation time influence the biological system: what forces do insects have to produce, and how much energy does it cost them to extract themselves from sticky PF compared to water?

Another property of PF that has been pursued in previous studies is the fluid’s surface tension (ST). Several studies have reported that insects sink more readily in PF than in water [16,18–20,23,24]. In Sarraceniaceae, ants sank rapidly in *Heliamphora sp.* fluid yet floated on rainwater [19], and in *D. californica*, 100% of the tested ants were retained while none broke the surface of pure water [20]. Additionally, an oiled needle repeatedly floated on water despite vigorous perturbation, while it readily sank in *S. flava* PF [18]. Quantitative ST measurements support these observations: fluids from open pitchers of *S. flava* and *D. californica* both produced ST values lower than water (66 mN/m and 47.9 mN/m, respectively) [18,20]. These findings confirm that ST is reduced in Sarraceniaceae PF, producing an air-fluid interface that is easier to penetrate than water. This fluid property can help explain the ‘sinking ants’ phenomenon: an insect falling into PF will mostly land on the fluid surface, but is then increasingly wetted through its struggles to escape, and sink [17,20]. The bacterial community in *D. californica* PF plays a role in reducing the ST, but it is unclear if the plants can also secrete surface-active compounds [20]. Nevertheless, these studies illustrate the importance of reduced ST for the effective retention of prey in Sarraceniaceae.

Meanwhile, the role of ST in *Nepenthes* remains the subject of debate. On one hand, there are several reports of ants readily sinking in *Nepenthes* PF: in *N. hemsleyana*, up to 80% of tested ants were completely submerged, compared to 10% in water [17]. Similar observations have been reported elsewhere [23]. However, ST measurements to date show that fluids from two *Nepenthes* species have ST close to that of water (72 mN/m for *N. rafflesiana* [13,17], 73 mN/m for *N. hemsleyana* [17]; 72 mN/m for water [13,17]). Hence, based on these contradictory findings, it is difficult to conclude if a reduced ST is responsible for sinking prey and if it influences insect retention in *Nepenthes* PF.

Here, we investigate the effect of sticky PF on insect retention, with a focus on adhesion of the fluid to insect cuticle. Using *N. rafflesiana* PF as our study system, we first quantify the forces exerted on an ant gaster (the abdomen) as it is wetted and then retracted from the fluid, thereby simulating an insect’s attempt to escape. Next, we re-assess the role of ST in prey retention through two distinct approaches. Lastly, we study the dewetting behaviour of PF on different surfaces and highlight a novel function of the viscoelastic nature of the fluid and a new mechanism of prey retention. Our findings illustrate the diversity of interactions between the fluid and the insect cuticle that becomes evident under dynamic testing conditions.

## 2. Materials and methods

### 2.1. Pitcher plant fluid samples

*N. rafflesiana* PF was collected from unopened pitchers that were close to opening in Brunei, northern Borneo (4°34’ N, 114°25’ E; collection site: degraded kerangas forest on white sandy soil) and from greenhouse cultivars at the University of Bristol (courtesy of Dr. Ulrike Bauer, University of Bristol; plants sourced from Brunei, Malesiana Tropicals nursery, or Kew Gardens). Each pitcher was either cut open with a clean razor blade and its contents poured into a sterile plastic collection vial (field collections), or its lid was opened and the fluid removed using a clean pipette (greenhouse collections). The samples were kept frozen at −20°C until use. Prior to experiments, individual vials were thawed to room temperature, and a small aliquot was stored in a 4°C refrigerator while the stock was re-frozen. PF was stored at 4°C for the duration of the experiments with no growth of contaminants or visible changes to the fluid consistency.

### 2.2. Observations of ant in pitcher plant fluid

Ants from a laboratory colony of *Atta cephalotes* were used to observe the behaviour of live insect prey in *N. rafflesiana* PF. Medium-sized worker ants were used (mean weight ± SD: 7.6 ± 1.3 mg). Ant behaviour was assessed using methods related to previous studies [13,16,17]: each ant was placed inside a slippery Fluon-coated container that was held 5 cm above the fluid surface (size of aquarium with fluid: 7.6 x 2.5 x 2.5 cm). The container was then slowly tipped so that the ant slid down the wall and into the fluid without manipulation or coerced acceleration. This method aimed to mimic ants naturally slipping on the peristome surface and free-falling into the pitcher. The ant’s behaviour was observed for 5 min and categorised using a metric derived from [16]: 1. ‘Escaped,’ if it removed all 6 legs out of the fluid; 2. ‘Walking on water,’ if its legs did not break the fluid meniscus and instead tried to walk on the surface; 3. ‘Swimming,’ if it tried to swim but without all of its body submerged below the surface; 4. ‘Floating motionless,’ if it stopped moving within 5 min but did not sink; ‘Sunken,’ if all of its body was completely submerged below the fluid surface. Ants were tested in sticky PF and in reverse-osmosis water (RO water; n=10 ants per fluid).

### 2.3. Force measurements on ant gasters

Ants were used to assess the range of forces experienced by insects that had fallen into sticky PF. Rather than using whole ants, which would introduce many uncontrollable variables regarding surface topography, orientation, and shape, we opted to use the gasters of similarly-sized ants as a model cuticle surface. Gasters were prepared as follows: an ant was euthanised by freezing, weighed, and its gaster cut at the petiole. Next, an insect pin was inserted into the cut end (see Supplementary Fig. 1. A small droplet of high viscosity cyanoacrylate adhesive was applied at the junction to immobilise the gaster. Special care was taken to avoid contaminating the cuticle with excess adhesive. Each mounted gaster was inspected under a stereomicroscope for contamination or damage prior to use.

A custom fibre-optic force transducer was set up as previously described with some modifications (Supplementary Fig. 1) [25]. A piece of reflective metal foil was glued onto a thin metal beam (beam spring constant: 2.5 N/m), and then mounted to the Z-motor stage of a 3D motor system (M-126PD, Physik Instrumente, Karlsruhe, Germany). The fibre optic sensor (DMS-D12, Philtec, Inc., MD, USA) was lowered towards the foil until the output was in the linear regime of the signal-to-distance curve. A signal-to-force calibration was obtained using pre-weighed objects and a signal-to-displacement calibration was calculated by incrementally bending the beam using the Z-motor. All motor movements and constant force feedback protocols were performed using LabVIEW (National Instruments, TX, USA). A USB camera synchronised to the motor movements was used to film the sample interacting with the test fluid (20 fps; DMK23UP1300, The Imaging Source Europe GmbH, Bremen, Germany).

A small aquarium with two contiguous chambers to hold the test fluids was 3D-printed (Zortrax Inkspire, Zortrax S.A., Olsztyn, Poland; Supplementary Fig. 1). This design allowed for the same specimen to be tested in two different fluids without re-mounting. One wall of the chamber was made with a glass coverslip to provide a clear view of the sample while filming. For each trial, the aquarium was rinsed once each with ethanol and RO water, then around 600 μL of PF or water was transferred into the two chambers. The insect pin (with a gaster attached) was bent so that the dorsal side of the gaster would be the first to contact the fluid. This reduced the likelihood of the stinger and abdominal glands interfering with the measurements (*e.g*., a sharp edge can pierce the meniscus, and dried glandular secretions can be hydrophilic). The sample was then attached to the force transducer using dental wax (Elite HD+, Zhermack SpA, Italy).

Once the gaster and the fluids were prepared, the following motor movements were executed: 1. Lower sample into the fluid to reach a preload of 50 μN (approximately two-thirds of a tested ant’s body weight); 2. Stay in force feedback-controlled preload for 4 s; 3. Move up by 10.5 mm at 3 mm/s; 4. Dry for 60 s with a small fan; 5. Repeat steps 1-3 twice to produce dip 1, 2, and 3. This protocol was designed to simulate repeated escape attempts by a fallen insect. The trials were paired so that each gaster was first tested in water (total of three dips), dried for at least 3 min, then tested again in PF. We conducted control experiments with water in both wells (water-water) to confirm that using the same gaster twice did not affect its properties (no significant difference in peak force between the first and second water tests; paired *t*-tests for dips 1, 2, 3: t_5_= −0.499, t_5_= −0.752, t_5_= −0.728, all *p* > 0.05; n=6 gasters).

From each gaster trial, we calculated the ‘work of retraction’ using the following equation:

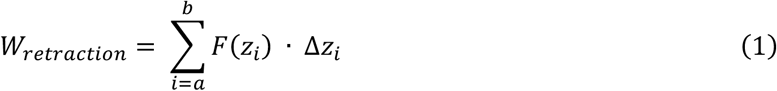

where *F*(*z*_*i*_) is the force measured at a specific motor position *z*_*i*_, Δ*z*_*i*_ is the distance moved between two force measurements, and *a* and *b* denote the start and end indices for the summation (see supplementary materials for details). The peak attractive force was defined as the maximum force recorded during the upward Z-motor movement. Six sticky *N. rafflesiana* PF samples were selected to measure the peak attractive force and work of retraction.

### 2.4. Statistical analyses

Restricted maximum-likelihood linear mixed-effects modelling was used to analyse the relationship between the dependent variables (work of retraction and peak attraction force; the former was natural log-transformed) and the independent variable (test liquid, with water and PF as levels, and dips). Test liquid and the interaction between test liquids and dips were used as fixed effect terms. Individual ant gasters and PF samples were used as random intercepts to account for the nested design and repeated sampling (each ant gaster tested first in water then in PF; six gasters tested per PF sample). Data from each dip was separated and analysed with the same parameters. *t*-tests were conducted via Satterthwaite’s degree of freedom method per the *lmerTest* package [26]. All tests were conducted in R v3.6.2 (run in RStudio v 1.2.5033) using R packages *lme4* v1.1-21 and *lmerTest* v3.1-1 [26–29].

### 2.5. Force measurements using antennae as model insect cuticle

Ant antennae are densely covered in hairs and preliminary tests confirmed that they strongly resisted wetting, and when submerged in water, a layer of air remained trapped between the hairs. Hence, antennae were used to test if PF could wet highly hydrophobic surfaces. Each antenna was cut at the end of the first segment and the cut end was attached to an insect pin with a small droplet of cyanoacrylate adhesive (inspection under a stereomicroscope showed that the adhesive did not spread on the antenna). Both antennae from each ant were mounted using the same technique to produce comparable samples. Extreme care was taken to avoid touching the last few segments of the antenna, and each sample was visually inspected for contamination or damage. Since freshly prepared antennae were highly flexible, all samples were stiffened by drying them in a desiccator for 2-3 hrs prior to use.

For each trial, one insect pin-mounted antenna was attached to the force transducer and positioned so that the last segment would be the first to contact the fluid surface. One of the two antennae from an ant (sample A) was used to determine the loading force at which the tip ruptured the RO water meniscus. The following protocol was then used to simulate an insect body-part falling onto the fluid interface and staying in contact for a period of time: 1. Slowly lower sample into the water until set preload; 2. Maintain preload using force-feedback for 60 s; 3. Return to the starting position. If the liquid meniscus did not break, then the loading force was increased by one 5 μN increment and the trial repeated (tested force range was 30-50 μN). The maximum preload was defined as the force which was one increment smaller than the force needed to rupture the meniscus. Control experiments showed an antenna could be tested 4-5 times at its maximum preload on water without a change in response (n=3 antennae), confirming that antennae were not wetted despite multiple dips. Nevertheless, to demonstrate that the antenna had not been wetted by water during the trials, every sample was tested twice at the same maximum loading force. If the antenna did not break the meniscus in both repetitions after 60 s, the trial was recorded as ‘meniscus held.’

Once the maximum loading force was determined, sample A was replaced with the second antenna from the ant (sample B) to test in PF. Due to random variation between the first and second antenna for the maximum sustainable preload force, we conservatively expected 50% of the tested antennae to break through and sink into the fluid. If the antenna penetrated the meniscus before the full 60 s preload, the sample was recorded as ‘meniscus broke.’ Six pairs of antennae were tested in each pair of water and PF (n=3 different PFs).

### 2.6. Measuring the surface tension of PF

Pendant drop tensiometry can be a highly accurate method of measuring surface tension with small volumes of test fluid if some precautions are taken to reduce experimental error [30]. The technique is based on analysing the shape of a static drop that results from a balance of its weight and the surface tension force exerted by a needle. This balance is represented by a dimensionless parameter called the Bond (or Eötvös) number:

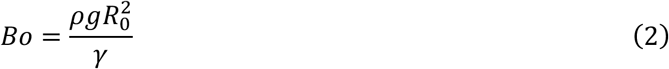

where *R*_0_ is the radius of curvature at the apex of the droplet, *γ* is the surface tension, and *ρ* is the density of drop. A second dimensionless parameter, the Worthington number, was incorporated in [30]:

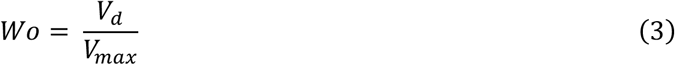

where *V*_*d*_ is droplet volume and *V*_*max*_ is the theoretical maximum volume that can be formed by the needle. When *W*o is greater than 0.6, then the experimental error is below 1%, hence it serves as a useful criterion for minimising error. We used a slightly modified version of the method described in [30]: ~50 μL of PF was withdrawn into a 1.0 mL syringe fitted with a blunt-ended needle (inner diameter: 0.8 mm). A syringe pump (AL-1000, World Precision Instruments, FL, USA) was used to dispense the fluid at a constant flowrate of 7 μL/min and filmed with a USB camera at 3 fps (Basler acA1300-200um, Basler AG, Ahrensburg, Germany). Ten PF samples from *N. rafflesiana* were tested. In addition, two PF samples from another species (*N. inermis* from Cambridge University Botanical Gardens) were measured as they were also sticky and viscoelastic. Between 5-7 drops with *W*o greater than 0.6 were selected for each sample and analysed using OpenDrop (http://opencolloids.com). All experiments were conducted at 25°C in ambient humidity (30-40% RH). The full dataset is provided in the supplementary materials.

ST measurements of distilled water were in good agreement with the reference value of 72.0 mN/m at 25°C [31]. To check if ST values of dilute viscoelastic fluids were in line with those obtained with the Du Noüy ring method [32,33], we tested solutions of commercial xanthan gum (Sigma-Aldrich; molecular weight: ~2×10^6^ Da; concentrations: 0.1, 0.2, 0.5% w/v in distilled water).

### 2.7. Visualising PF dewetting behaviour on various substrates

A number of gasters and antennae tested in water and PF were subsequently imaged with scanning electron microscopy (SEM; n=22 antennae, n=20 gasters). Unused gasters prepared under the same conditions served as controls. Specimens were mounted on aluminium stubs, coated in 15 nm of iridium, and imaged with an SEM (Verios 460, ThermoFisher Scientific, MA, USA).

To investigate how PF interacts with different surfaces, we recorded the dewetting of PF and water from three surfaces: clean glass coverslip (hydrophilic), clean smooth low-density polyethylene (PE, hydrophobic), and *A. cephalotes* gaster cuticle (freshly prepared as described above). Static contact angles of water on the glass and PE surfaces were measured using a goniometer (7 μL per droplet; n=10 droplets per surface; OCA 15EC, DataPhysics Instruments GmbH, Filderstadt, Germany). The static contact angles were 51.9° ± 0.4° and 86.3° ± 0.8° (mean ± standard error of the mean) for glass coverslip and PE surfaces, respectively. We used interference reflection microscopy (IRM; as reported previously in [34–36]) to visualise the formation and evolution of films during dewetting. More specifically, a small volume (~5 μL) was first deposited on the test surface using a micropipette. A clean microcapillary tube connected to a microinjector (CellTram Air, Eppendorf, Hamburg, Germany) was used to manually withdraw the fluid. The dewetting process was filmed at 25 fps (DMK23UP1300), and monochromatic green light was used for illumination (wavelength = 546 nm). Each fluid (n=2 different PF samples; RO water) was tested 3 times on glass, PE, and insect cuticle. PE surfaces were imaged using SEM after PF dewetting trials.

## 3. Results

### 3.1. Ant retention rates and behaviour in PF compared with water

Our retention trials with ants dropped in *N. rafflesiana* PF compared to water revealed a striking difference in outcome (Fig. 1a & b; Supplementary Video 1 and caption in supplementary materials). While none of the ants dropped in water sank and 30% ‘walked’ on the water surface without breaking the meniscus, all the ants dropped in PF were wetted upon landing and none managed to ‘walk’ on the PF meniscus (Fig. 1c). Moreover, 20% of the ants were fully submerged and sank within 5 minutes in PF. Ultimately, none of the ants managed to escape from PF, as opposed to 30% in water. Comparable observations and results have been previously reported for *N. rafflesiana* and *N. hemsleyana* using different species of ants [17].

**Fig. 1.**
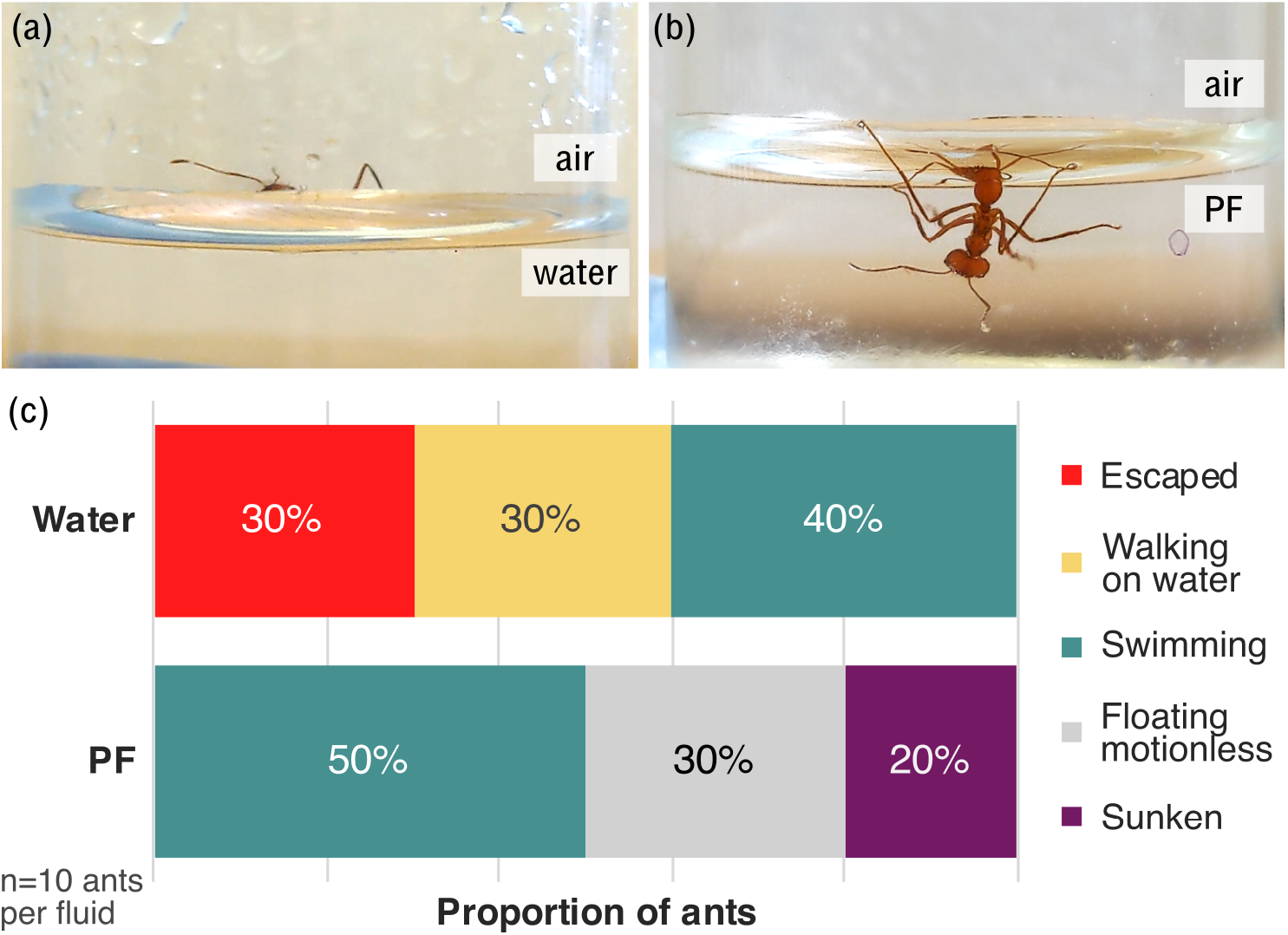
Ant retention tests in water and *N. rafflesiana* pitcher fluid (PF). (a) During retention tests in water, ants (*Atta cephalotes*) sometimes floated and failed to break the water meniscus. (b) When dropped into *N. rafflesiana* PF, ants readily broke through the pitcher fluid meniscus. Here, the test subject was submerged and failed to right itself. (c) The behaviour of the test ants was recorded for 5 minutes. While 30% of the ants escaped from water, none escaped from pitcher fluid. Additionally, 20% of the ants sank into pitcher fluid, which was not observed with water; instead, 30% walked on the water meniscus.

### 3.2. Force and energy required to retract ant cuticle from PF in simulated escapes

Using ant gasters mounted on the force transducer set-up, we simulated the movements of an insect attempting to escape from PF or water and measured the forces (see Fig. 2 for a representative force-time plot). As shown by the jump into contact when the gaster approached the water surface (Fig. 2a), all tested gasters were wetted from the first dip. Although insect cuticle is covered by a waxy layer, it is known that cuticle wettability depends on the species, specific body-part, and surface roughness (from hairs) [37–39]. Nevertheless, *A. cephalotes* gasters reached the desired preload of 50 μN with a partial immersion (Fig. 3a-1). Upon retraction, a liquid bridge formed between the water meniscus and the gaster and then rapidly collapsed, resulting in a sharp attractive force peak (Fig. 3a-2 & 2b-2). A small water droplet remained on the gaster immediately after the liquid bridge collapsed, but it often evaporated before the next dip. SEM images of samples tested in water showed no signs of contamination or residues (see Section 3.5).

**Fig. 2.**
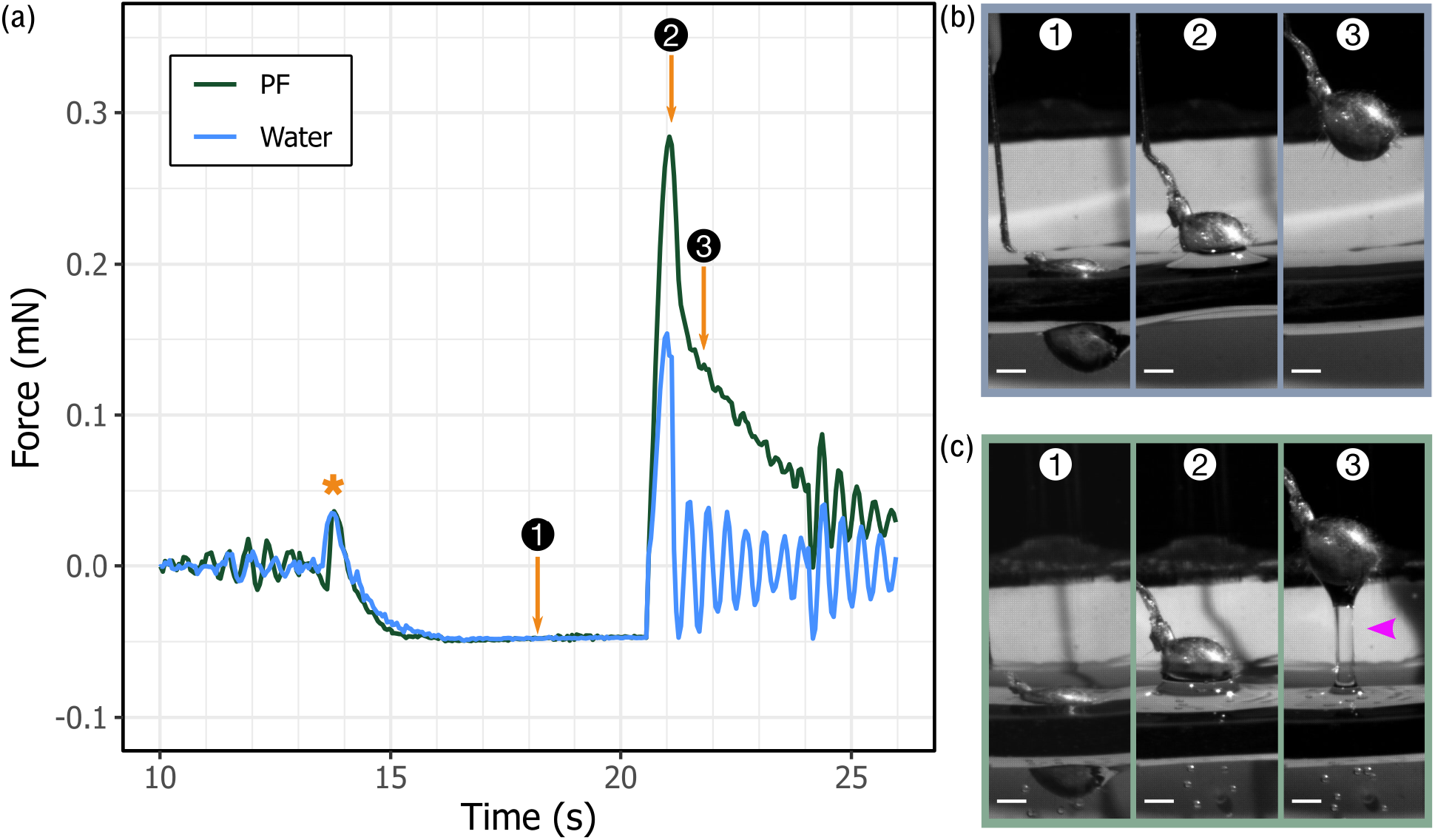
Representative force-time plot from force measurement trials of ant gasters dipped into water compared to *N. rafflesiana* pitcher fluid. (a) The gaster was lowered into the test liquid with 50 μN preload force, maintained for 4 s, then retracted upwards (note: a positive force is attractive). A small force peak resulted from the gaster jumping into contact (marked *). ①, ②, and ③ correspond to time-points 18.2 s, 21.1 s, and 21.8 s, respectively. (b-1 to 3) Images taken at the time points ① to ③ as shown on the force plot. At ①, the abdomen was preloaded. Upon withdrawal from water, a liquid bridge between the abdomen and the fluid rapidly collapsed (②), creating a sharp attractive force peak (③). (c-1 to 3) In contrast, when the gaster was withdrawn from pitcher fluid, a liquid bridge formed (③; arrowhead), resulting in a higher peak attractive force and a slower and prolonged decay of the attractive force. Scale bars for (b) & (c): 500 μm.

**Fig. 3.**
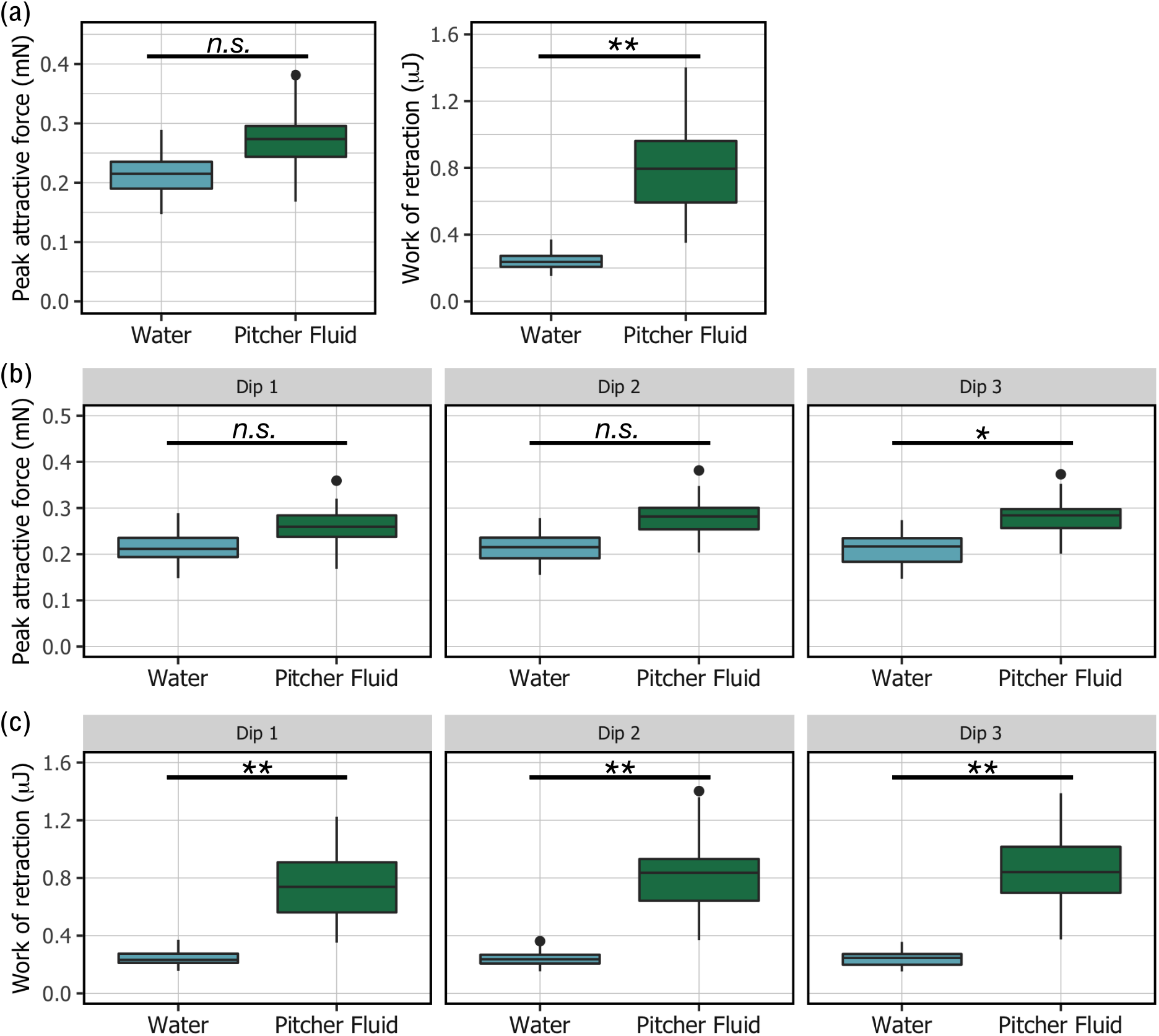
Effect of *N. rafflesiana* pitcher fluid (PF) on the peak attractive force and work of retraction for an ant gaster. (a) Overall, the peak attractive force acting on the ant gaster during retraction was marginally higher in PF than in water but not significantly (n.s., *p*=0.12). The work of retraction, however, was 2.9 times greater in PF than water (\*\**p* < 0.01). (b) When separated into the individual dips, PF exerted a significantly higher peak attractive force on ant gaster than water only by Dip 3 (\**p* < 0.05). (c) In contrast, PF consistently demanded higher work to retract within each dip compared to water (\*\**p* < 0.01). All statistical analyses were based on *t*-tests on linear mixed effects models (see text for details).

When gasters were preloaded and retracted from PF, however, the peak attractive force was marginally larger than in water (Fig. 2a). Moreover, upon retraction from PF, we observed filament formation between the cuticle and the fluid surface (Fig. 2c). The adhesive effect of the filament, which essentially pulled the gaster back into the fluid, was visible on the force-trace as a slower and prolonged decay of the peak force (Fig. 2a-3 & 2c-3). We also observed that after retracting the gaster from PF and the collapse of the liquid bridge, a much larger droplet formed on the gaster compared to the test in water. This became more pronounced after the 60 s of drying in air, where the water droplets were either significantly smaller or no longer visible, while a large PF droplet remained on the gaster.

Comparisons between the peak attractive forces from all water and PF trials indicated a trend for a greater attractive force in the latter, but this was not statistically significant (Fig. 4a; *t*-test based on the aforementioned linear mixed effects model, t_5.1_=1.857, *p=*0.12). In contrast, the work required during the simulated escape movement in PF was 2.9 times greater than in water (*t-*test, t_5.0_=5.590, *p* < 0.01), caused by the sustained adhesive force from the PF filament.

**Fig. 4.**
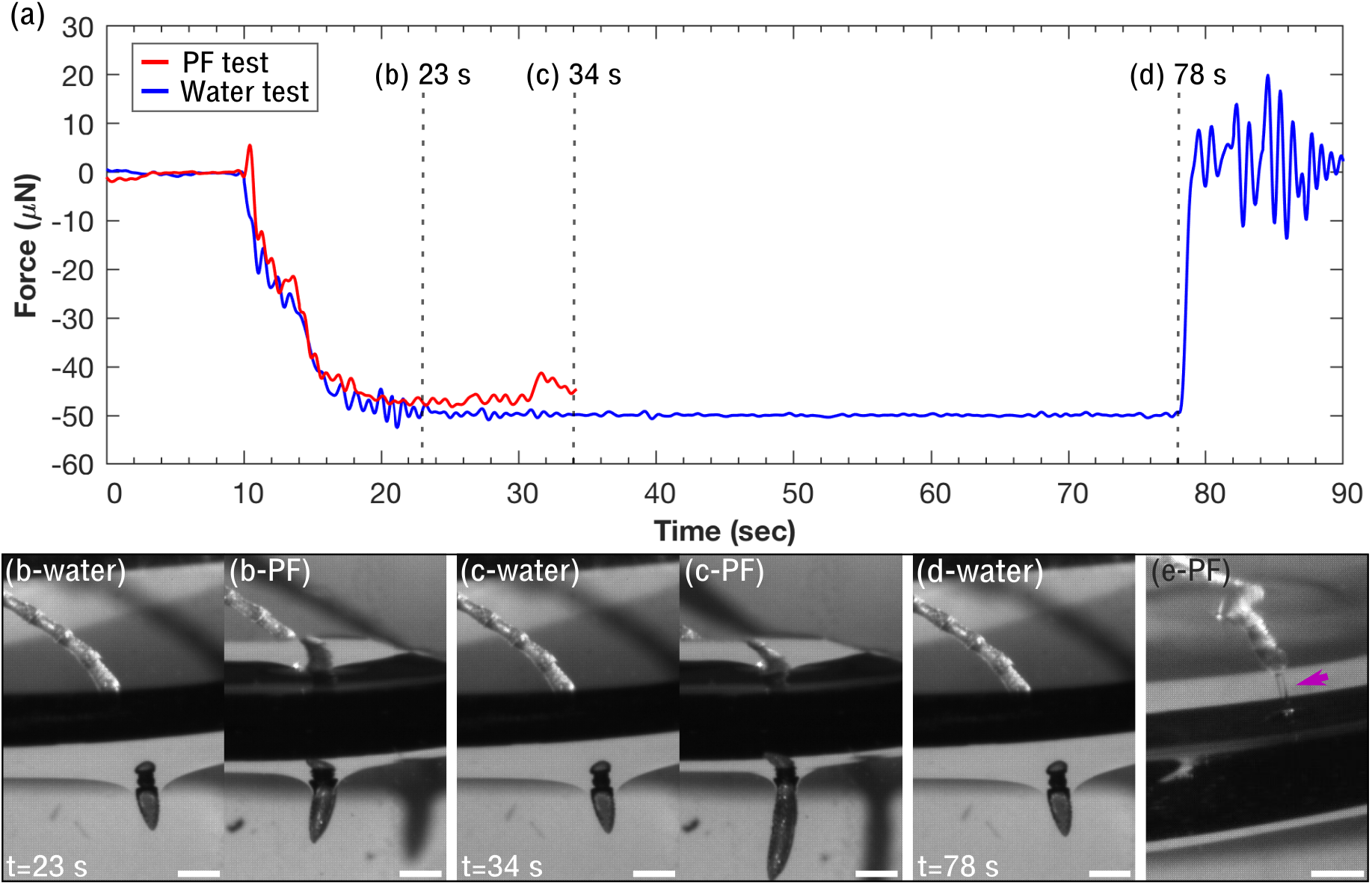
Using ant antennae to probe the surface tension of water versus pitcher fluid. (a) The ant antenna did not break through the water meniscus at the designated preload force (50 μN in this example) for the entire duration of the trial. The other antenna from the same ant tested in pitcher fluid failed to reach the designated preload as it readily broke through the meniscus and the movement was terminated. (b & c) Image sequences highlight the difference between the water and pitcher fluid trials. From 23 s to 34 s, the water-test antenna held steady at 50 μN preload, while the pitcher fluid antenna was pushed deeper into the fluid as it failed to reach the preload. (d) Over the full duration of the trial, the water meniscus remained steady. (e) A fluid filament formed upon withdrawal of the antenna (see arrow). Scale bars: 500 μm.

When we analysed the effect of dips on the peak attractive force, we found that within each dip, the peak attractive force for PF was marginally but significantly higher than water by the third dip (Fig. 3b; *t-*test, t_2.0_=4.305, *p* < 0.05). Across the dips overall, we found a significant interaction between the dips and the peak attractive force, where the effect of dips on peak attractive forces significantly depended on the tested fluid (*t-*test, t_178_=3.455, *p* < 0.001). Subsequently, when water and PF were analysed separately, we confirmed that dips had no significant effect on the peak attractive force in water (*t-*test, t_74_=−0.55, *p* =0.584), while it had a clear impact on PF (*t-*test, t_74_=5.243, *p* < 0.001). Thus, peak attractive force did not change from multiple dips in water, whereas there was an increase in PF.

In contrast to the peak attractive force, the work of retraction required to withdraw gasters from PF was consistently higher than in water for all three dips (Fig. 3c; *t-*test, Dip1: t_2.59_=7.071, *p* < 0.01; Dip2: t_3.09_=7.334, *p* < 0.01; Dip3: t_3.55_=6.904, *p* < 0.01). The overall findings for the effect of dips on the work of retraction were similar to those stated above for the peak attractive force. Implications of these results are addressed in the Discussion.

### 3.3. Loading the liquid-air interface to test if PF meniscus breaks more readily than water

From our experiments using antennae to probe the liquid-air interface, we first identified the maximum preload force that the water meniscus could sustain (sample A, n = 16 antennae; Fig. 5a). All of these samples failed to break the water-air interface to advance into the fluid during the entire 60 s preload (Fig. 5b-d). Video recordings showed that while the tip of the antenna broke through the meniscus, the water contact line was arrested at or before the widest point of the antenna and held for 60 s. For antennae (sample B) dipped in PF, however, the outcome was clearly different: 15 of the 16 antennae (93%) broke through the meniscus and continued to advance within 60 s (Fig. 5b-PF & 1c-PF). Even using the most conservative assumption that 50% of the antennae break the meniscus by chance, this result is highly significant (binomial test; *p*=2.59×10^−4^). Furthermore, of the 15 antennae that broke through the meniscus, four formed thin filaments upon withdrawal from the PF (Fig. 5e-PF).

**Fig. 5.**
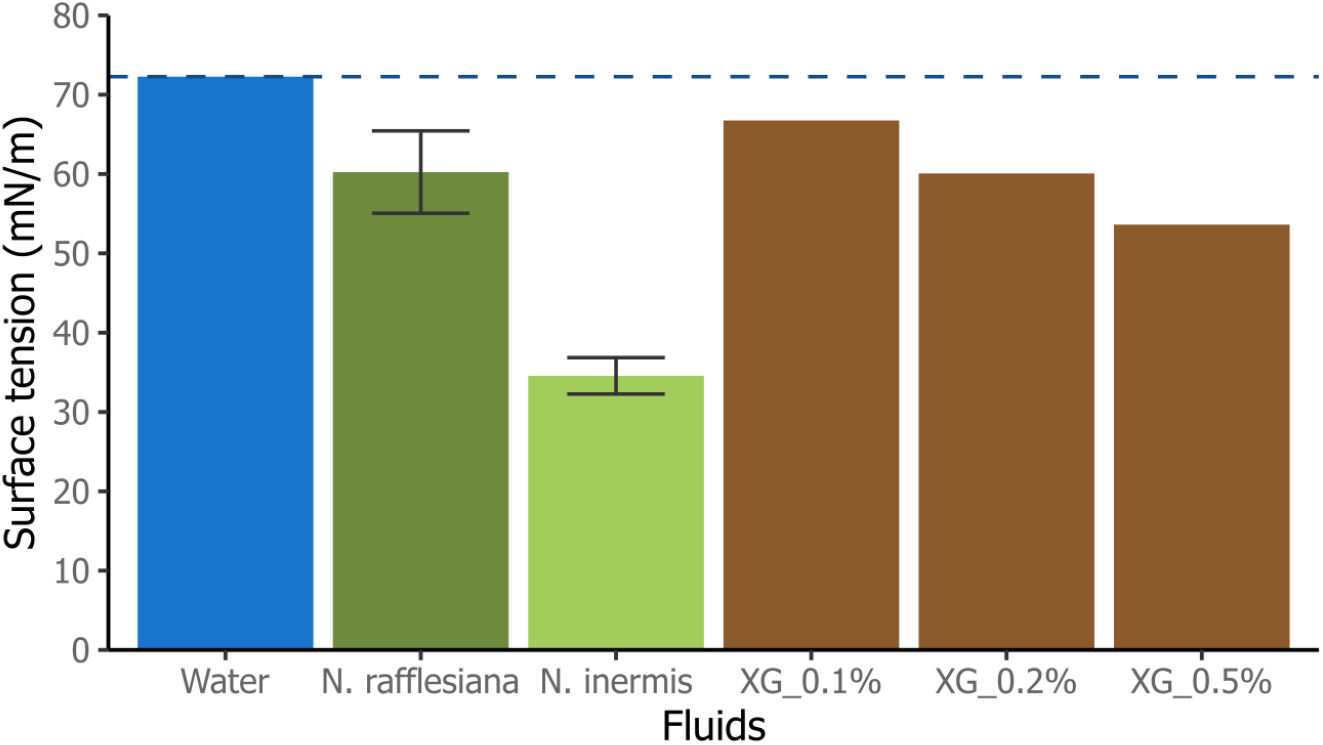
Surface tension values of *N. rafflesiana* and *N. inermis* pitcher fluids compared to standard fluids as measured by pendant drop tensiometry. Good agreement between measured surface tension of water and reference value at 25°C validated the method (72.3 ± 0.6 mN/m and 72.0 ± 0.36 mN/m, respectively). *N. rafflesiana* pitcher fluid surface tension values were significantly lower than reference value of water (*t*-test, t_9_=−7.13, *p* < 0.001). *N. inermis* pitcher fluid produced the lowest surface tension value out of all tested fluids. Error bars shown only for *N. rafflesiana* and *N. inermis* (± standard deviation; see main text for water and xanthan gum values). Increasing concentrations of commercial xanthan gum (XG; w/v) led to a decrease in surface tension, a trend reported in other studies.

To confirm that our findings were not affected by variation from switching sample A to B (for e.g., mounting orientation), we tested the same antenna without remounting: first in water, then in PF (n=12). None of the tested antennae broke through the water meniscus, even after two to five repeats; in contrast, 92% (11 antennae) broke through the PF meniscus. Again, even when the most conservative assumption was used (50% of the antennae break the meniscus by chance), this result is highly significant (binomial test; *p*=0.0032).

### 3.4. Surface tension of *Nepenthes* PF and water

ST measurements using pendant drop tensiometry further substantiated our finding that ST is reduced in sticky *N. rafflesiana* PF (Fig. 6). The ST value for water did not differ significantly from the reference value, which confirmed the validity of our method (72.3 ± 0.6 mN/m, mean ± SD; n= 10 droplets; one-sample *t*-test, t_9_=1.44, *p* = 0.18). On the other hand, the ST of *N. rafflesiana* PF was significantly lower than that of water (60.2 ± 5.2 mN/m, mean of means ± SD of means; n=10; one-sample *t*-test against reference value for water, t_9_=−7.13, *p* < 0.001). Preliminary tests from *N. inermis* PF produced a ST value of 34.6 ± 2.3 mN/m (n=2). The ST of solutions of commercial xanthan gum were (mean ± SD): 66.8 ± 0.1 mN/m, 60.1 ± 0.5 mN/m, and 53.6 ± 0.7 mN/m, for concentrations 0.1, 0.2, and 0.5% w/v, respectively. These values generally agreed with literature values and followed the same trend, where an increase in xanthan gum concentration led to a decrease in ST [32,33].

**Fig. 6.**
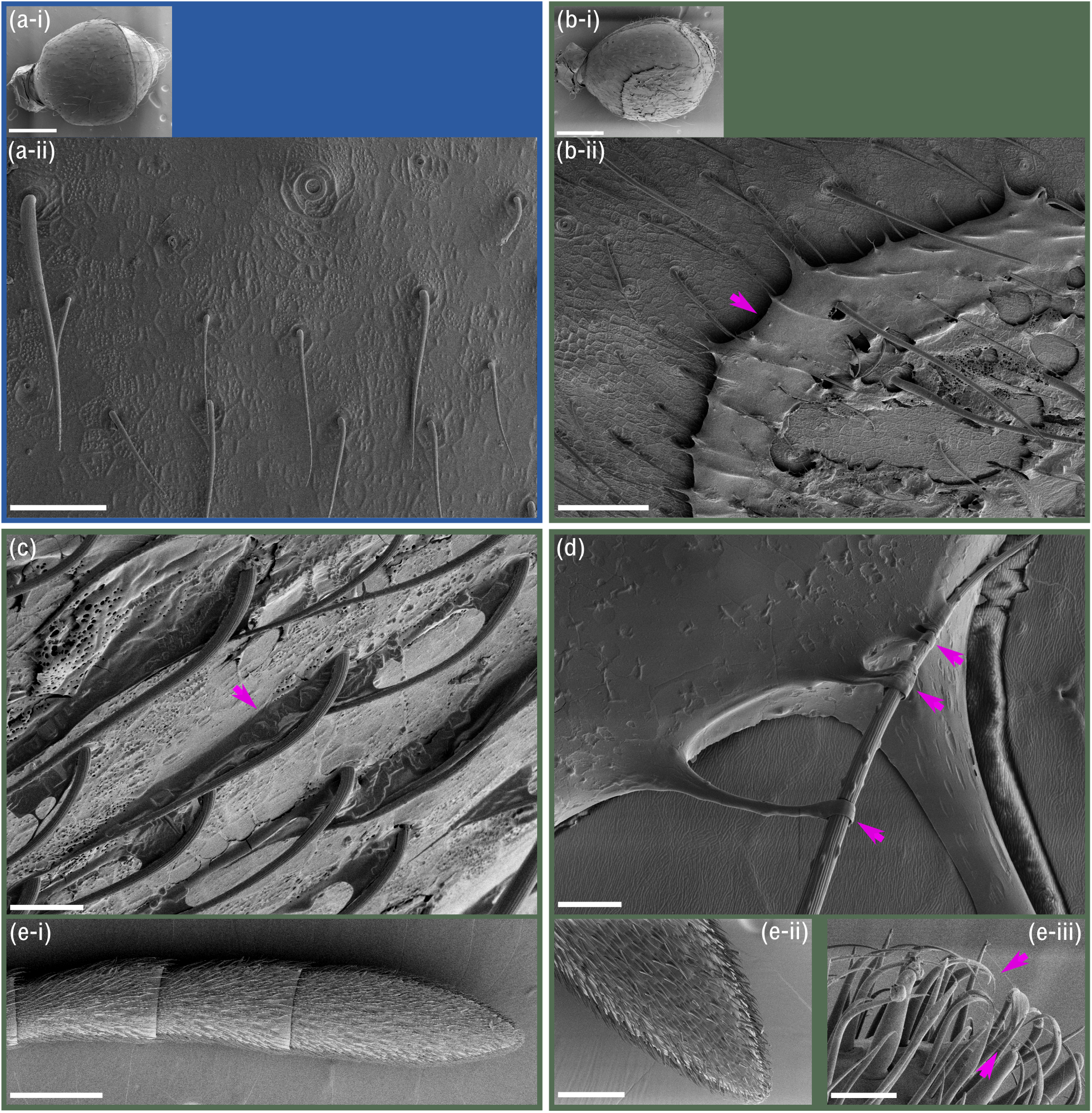
Scanning electron microscopy (SEM) images of ant gasters after testing in water (a, blue frame) and pitcher fluid (b-e, green frame). (a-i & ii): Gasters tested in water had no visible contaminants or residues on their cuticular surfaces. Scale bars: (a-i) 500 μm; (a-ii) 50 μm. (b-i & ii) Large areas of the gasters tested in pitcher fluid were covered by solid films of dried pitcher fluid (see arrow), coating both hairs and the cuticular surface. Scale bars: (a-i) 500 μm; (a-ii) 50 μm. (c) Dried pitcher fluid bridges between hairs and the cuticular surface (see arrow). Scale bar 20 μm. (d) Filaments ‘gripping’ a single hair (each filament marked by arrow). Scale bar 5 μm. (e-i) Last three segments of an ant antenna. The entire antenna is densely covered in sensory hairs. (e-ii) Antennae were generally less contaminated with pitcher fluid residues than abdomens. (e-iii) Closer inspection of the antenna tip revealed pitcher fluid filaments between the hair tips (but not the cuticle between the hairs; see arrows). Scale bars: (e-i) 250 μm; (e-ii) 100 μm; (e-iii) 10 μm.

### 3.5. Conspicuous residues are present on insect cuticle after contact with PF

After trials in PF, gaster cuticle and hairs were clearly coated in residues (Fig. 7). We observed films on significant areas of the gaster (smooth texture in some areas, porous in others; Fig. 7b-i & ii). Dried liquid bridges between the hair and the gaster cuticle were also visible (Fig. 7c). Polygonal crystals were sometimes present on the smooth cuticular surface. We also observed PF ‘gripping’ individual hairs to form solid filaments spanning between the main film and the hairs (Fig. 7d). Note that these samples were not flash-frozen and freeze-dried but rather dried in a desiccator over several days, hence the filaments were stable structures. Gasters tested in water were free of any visible residues and closely resembled the control samples (Fig. 7a-i & ii).

**Fig. 7.**
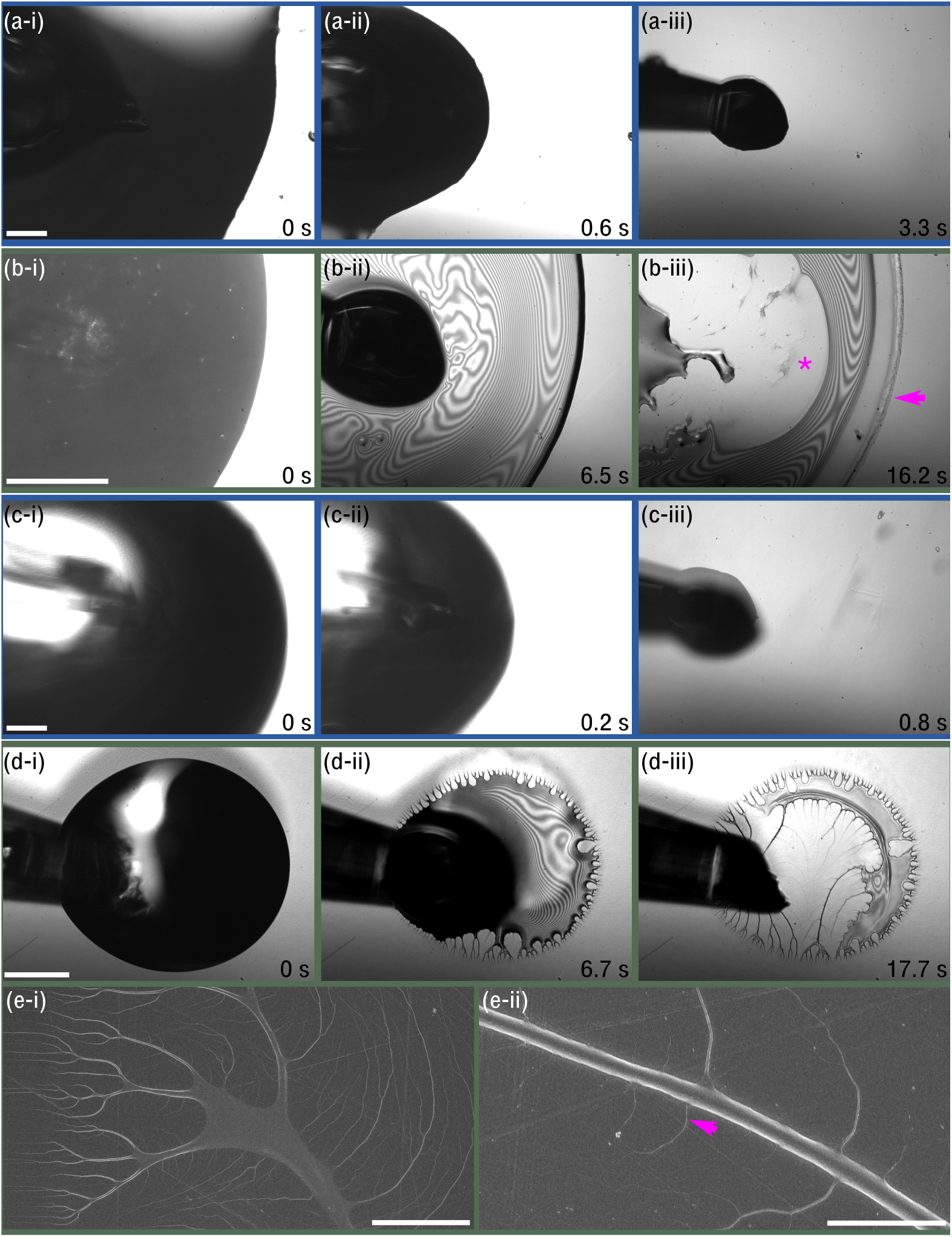
Dynamic dewetting behaviour of water (a,c - blue background) and *N. rafflesiana* PF (b,d - green background) and on different surfaces visualised via interference reflection and scanning electron microscopy. (a-i, ii, iii) Water droplet retracted from clean glass (hydrophilic) surface. The droplet dewetted cleanly without leaving residues within 3.3 seconds. (b-i, ii, iii). In contrast, PF resisted dewetting from glass, as shown by the formation of a thin layer (interference fringes visible in ii). Even after 16.2 s, there were residues on the surface, and the initial outermost rim had not contracted (arrow). Very thin films were left behind (marked by *). (c-i, ii, iii) On polyethylene (PE, hydrophobic) surfaces, water droplets completely dewetted. (d-i, ii, iii) PF, on the other hand, behaved similarly as on glass, where a thin layer was formed as more liquid was withdrawn. Moreover, solid fractal-like filaments were deposited on the surface. By 17.7 s, the surface remained partly coated by dried PF films and filaments. (e-i, ii) SEM of the fractal-like filaments on PE surface. Extremely fine filaments (~20 nm in diameter, see arrow) were also present on the surface. Scale bars: (a-d) 200 μm; (e-i) 20 μm; (e-ii) 2 μm.

Assessment of tested ant antennae using SEM highlighted the high density of cuticular hairs on the antennae, which inhibited wetting of the underlying smooth cuticle. We observed that antennae tested in water were mostly free of filaments (Fig. 6e-i & ii), although dirt-like particles were present at the tip of some specimens (n=4 out of 7 antennae). The majority of the antennae tested in PF were free of residues, but in some cases we observed filaments spanning between the hairs at the tip of the antennae (Fig. 6e-iii; n=4 out of 15 antennae). No other residues or contaminants were found on the remaining antennal segments, and overall the antennae were cleaner than gasters after tests in PF.

### 3.6. Dewetting is slowed down or prevented in PF

Water dewetted from the clean glass surface without leaving behind any residues or films (Fig. 8a-i to iii; Supplementary video 2). In stark contrast, PF on glass did not show any dewetting: the initial perimeter of the droplet did not contract, and continued withdrawal led to an increasingly thin film (Fig. 8b-i to iii; Supplementary video 3). Eventually, the film started to dry close to the microcapillary tube, and then at the outer fluid perimeter. This resulted in the formation of very thin layers or smaller filaments on the surface (see asterisk in Fig. 8b-iii), although it is possible that the PF dewetted in between these regions. When the microcapillary was removed and the fluid began to evaporate, branched hygroscopic crystals formed that visibly absorbed the humidity from our breaths when we blew on the glass surface (Supplementary video 3). On hydrophobic PE surfaces, water also dewetted completely from the surface (Fig. 7c-i to iii). In contrast, PF formed thin layers and residues on PE surfaces (Fig. 7d-i to iii). Furthermore, we observed fractal-like patterns comprised of solid filaments extending from the edge of the initial rim towards the centre (Fig. 7d-iii). SEM images of the PE surfaces confirmed that PF does not completely dewet from the surface (Fig. 7e-i & ii): we observed fine filaments coating the surface, and while regions between the micrometre-scale filaments looked clean, higher magnification images revealed numerous filaments with diameters less than 100 nm.

On insect cuticle, small droplets of water readily dewetted and evaporated from both smooth cuticle and cuticle with large hairs (Supplementary video 4). PF, however, behaved differently on cuticular surfaces: although the strong reflectivity of the cuticle obscured any interference patterns during the withdrawal, PF again failed to fully dewet from the cuticle, leaving residues on some areas of the cuticle, similar to those on glass and PE surfaces (Supplementary Fig. 2. The residues also resembled the patterns visible on the SEM images of cuticle after PF tests (Fig. 6b-i). Moreover, we observed filaments forming between the receding PF and hairs on the gaster, which could result in the aforementioned ‘hair-gripping’ structures (Fig. 6d).

## 4. Discussion

Carnivorous plants have evolved a myriad of adaptations and mechanisms to prey on insects, ranging from trigger hair-activated leaves of Venus fly-traps, sticky ‘glue’ secretions of sundew plants, and pitfall traps of pitcher plants. Although there are several structural adaptations that facilitate insect capture and retention in pitcher plants, it is increasingly evident that the digestive fluid itself can contribute mechanically to the capture and retention of insect prey. In the case of *N. rafflesiana* pitcher plants, one of several *Nepenthes* species that produces sticky PF, previous researchers have proposed two mechanisms responsible for the higher prey retention rate of the fluid compared to water. First, insects that fall into the fluid struggle and move rapidly to their own detriment: the viscoelastic shear-thinning fluid may respond elastically to fast shear-rates, which is thought to inhibit movement and hamper escape [13]. Second, as the insect retracts its wetted limbs from the fluid during its struggles, filaments are formed that resist being stretched, a consequence of the fluid’s high extensional viscosity [13]. For both of these mechanisms to work, we need to assume that the fallen insect is wetted by the fluid. Insect cuticle, however, is covered by cuticular hydrocarbons [37,40–42], which (often in combination with cuticular micro- and nanostructures) makes it generally difficult to wet with water. Moreover, small insects like ants and flies have low mass while the surface tension of water is high, which means they will not easily break through the meniscus to become submerged in the first place. Our work on the interactions between PF and insect cuticle provide insights into the mechanisms underlying the adhesive and retentive property of sticky PF, which are discussed further below.

### 4.1. Lower surface tension of PF facilitates sinking of insect prey

By dropping ants into PF to mimic natural insect capture events, we found that ants sank in PF but not in water. Additionally, ants ‘walked’ on the water surface, which was not observed in PF. This suggests that ants break through the PF-air interface more easily than the water-air interface, which is consistent with previous reports of insects sinking in *N. rafflesiana* fluid [17,23]. Using an antenna, a highly non-wetting part of the ants’ cuticle, we confirmed that PF produced a smaller up-thrust than water. We found that at the maximum preload force (41 μN on average, corresponding to approximately half the average weight of the tested ants), the up-thrust in water prevented the antenna from sinking in deeper. In PF, however, the antenna was pushed further into the fluid.

This result can be explained by a lower surface tension of PF compared to water. The upward force on an object at the fluid-air interface depends on the ST of the fluid and the surface wettability of the object, but the relative contribution of both factors is a function of the geometry of the object [43,44]. If we consider the simple case of a long and smooth cylinder with its axis perpendicular to the liquid-air interface, the ST force *F*_*w*_ is:

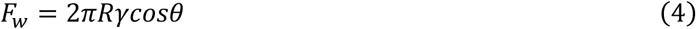

where *γ* is ST, *θ* is the contact angle, and *R* is the radius of the cylinder. A negative *F*_*w*_ corresponds to an up-thrust. Using Young’s law:

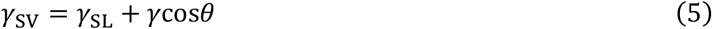

the equation can be rewritten as:

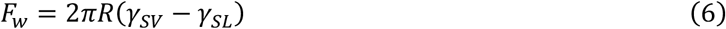

where *γ*_SV_ and *γ*_SL_ are the solid-vapor and solid-liquid interfacial tensions. Equation 6 predicts that *F*_*w*_ does not depend on the ST itself but only on the wetting via *γ*_*SL*_; higher *γ*_*SL*_ values (corresponding to larger contact angles for a given value of *γ*) would result in more negative values of *F*_*w*_ and hence more up-thrust. In the case of the antenna, however, there is a stable layer of air trapped under the antennal hairs (in both PF and water), such that the Cassie-Baxter equation of wetting applies [45]:

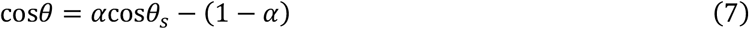

where *α* is the fraction of the surface occupied by a solid with contact angle *θ*_*s*_ and (1-*α*) is the fraction occupied by air. Combining Eq. 4, 5, and 7 gives:

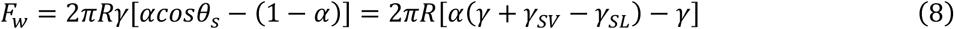

For the antenna, which has a dense covering of thin hairs, the solid area fraction α may be very small, so that:

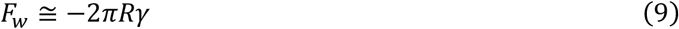

implying that the force is mainly dependent on ST (and less on wetting). A further geometrical argument supporting the importance of ST for floating objects can be made when considering a small horizontal cylinder floating on the liquid-air interface with its axis horizontal. Here, the dependence on the contact angle θ (provided that θ > 90°) is predicted to be weak [43,46] so that again ST would dominate. Therefore, the inability of the PF meniscus around the antenna to withstand the same force must arise from a reduced ST. Indeed, our pendant drop tests confirmed that sticky PF from both *N. rafflesiana* and *N. inermis* have lower ST than water. Thus, our force and ST measurements provide new evidence that reduced ST is important for prey capture and retention in *Nepenthes* PF, a mechanism long suspected but unsubstantiated until now [23].

We acknowledge that our ST values differ from previously reported values from the same species (*N. rafflesiana*), and some possible explanations are presented. One study used the capillary rise method, where the vertical rise of PF from unopened pitchers was measured in the field with 10 μL microcapillary tubes [17]. Given the small volume of the capillary and the high accuracy of the method under ideal conditions, it is unlikely that a ~15% change in ST (and thus the height) would have been missed. Instead, the discrepancy between our values may result either from experimental conditions or biological variation.

Another study used the pendant drop technique with PF from opening *N. rafflesiana* pitchers and found no significant difference to water [13]. One possible explanation is experimental error from how the droplets were dispensed: according to the study methods, a Pasteur pipette was used to form the droplets, implying that droplets were dispensed manually. Vibrations from manual injections could cause the droplet to pinch off prior to the maximum droplet size, which is a known source of error in pendant drop tensiometry [30]. Furthermore, at large pipette diameters and low Bond numbers (*e.g*., when the deviation from sphericity is small, as seen in fluids with surface tension close to water), there can be significant variation in the measured ST values, ranging from ~60 to ~95 mN/m for water [30]. To address this source of error, researchers used the Worthington number (*W*o) [30]. The study found that *W*o greater than 0.6 corresponded to a standard error of less than 1%; thus, we ensured that the *W*o value for each of our droplet measurements was greater than 0.6 (Supplementary Table 1). While the sample size was large and the variation in ST was low in the aforementioned study, Bond numbers were not reported [13]. Given the viscoelastic nature of PF, it is important to conduct each measurement at sufficient Bond or *W*o numbers to minimise variation. It is worth highlighting that *N. inermis* PF had a ST of 34.6 mN/m, which is nearly half of the ST of *N. rafflesiana*, and is lower than any of the previously reported measurements from Sarraceniaceae [18–20,23]. These findings further substantiate the idea that a reduced ST is a natural property of PF in some *Nepenthes* species.

Only a few studies have attempted to identify the surface-active molecules responsible for the reduced ST in PF. Initial tests with several members of Sarraceniaceae failed to detect saponins [18], a type of organic surfactant found in various plants [47,48]. Recently, both the ant retention rate and the ST value of *D. californica* PF were reproduced by inoculating sterile growth media with bacteria from PF [20]. This suggests that the bacterial community within *D. californica* pitchers not only help to break down organic matter [49] but also help lower the ST and improve prey retention. It is unclear whether the ST reduction is a by-product of the bacteria, or if the bacteria or the plant actively secrete surfactants.

Likewise, little is known about the molecules responsible for the reduced ST of *Nepenthes* PF. Although bacteria can reduce the ST in *Sarracenia*, we do not know if they also influence the ST in *Nepenthes*. However, it is worth mentioning that, due to the physiochemical properties of *N. rafflesiana* PF, neither bacteria nor chemical surfactants may be necessary to lower the ST. Previous researchers failed to detect bacteria in PF from unopened pitchers of several *Nepenthes* species [50], which suggests that bacteria may not be involved. In addition, *N. rafflesiana* PF is acidic [16,51], and organic acids have lower ST than water [52,53]. For example, a 1.6% w/w solution of acetic acid has a ST of 61.7 mN/m at 25°C [52], hence, the acidic nature of *N. rafflesiana* PF may be sufficient. Furthermore, it has been hypothesised that large (high molecular weight) polysaccharides are present in *N. rafflesiana* PF [13,54]. This could be important since a dilute solution of large polysaccharides alone can lower the ST: guar gum, for example, has a ST of ~50 mN/m (0.8% w/v) [55], and xanthan gum of 42.3 mN/m (1% wt) [56]. Dilute solutions of mamaku gum, a large polysaccharide with a chemical structure similar to the polysaccharide component of *Drosera* mucilage [57,58], yield ST values of 33.5 to 44.6 mN/m depending on the concentration [59,60]. Finally, the combination of low pH and large polysaccharides can act synergistically to further reduce the ST: for xanthan gum, ST decreases from 67.1 mN/m to 63.4 mN/m at pH 5 and 2.5, respectively [61]. Since *N. rafflesiana* PF reaches pH values as low as 2 [51] and is thought to contain acidic polysaccharides related to those found in *Drosera* mucilage [13,54], these are likely important parameters that influence the ST of *Nepenthes* PF.

### 4.2. Insect cuticle adheres strongly to PF

Surface wettability of insect cuticle is influenced by the outer lipid layer as well as surface patterning, often in the form of dense arrays of hairs and/or cuticle microstructures [62]. The initial wetting of ant gasters during our trials is likely based on slightly hydrophilic cuticle regions with advancing contact angles <90°; higher attractive forces during pull-out may be based on surface roughness and chemical heterogeneity, which reduce the receding contact angle. A similar argument was used in a study that examined why water alone retained certain species of ants and flies but not others: those with more wettable cuticular surfaces may wet more easily and thus have higher likelihood of sinking [16]. Consequently, when ant gasters were tested in water, we observed an overall repulsive force during the preload phase and a sharp adhesion peak when retracted. The transient adhesive peak occurred just before the rapid collapse of the meniscus; hence, if a trapped ant is able to find a surface to adhere to and overcome this peak through a short yet forceful burst of movement, it could escape from water. As mentioned previously, pitfall traps likely address this with structural adaptations on their inner walls to prevent the insects from gaining a sufficient foot-hold. The situation is different with winged insects, however, as they can take-off and fly away from the surface; consequently, their retention rate in pure water is low [16]. In such cases, sticky PF offers a clear advantage over water: while the peak attractive force is only weakly higher than for water, the displacement and hence the work of retraction is significantly larger. This is likely due to two factors: (1) the meniscus between the fluid and the gaster acts to pull the latter back; (2) a droplet remains adhered to the gaster and resists dewetting, adding weight and also facilitating re-wetting of the cuticle (see below). Thus, insects have to sustain higher forces for longer durations to escape from PF compared to water. For nonflying prey like ants, the adhesion of PF to their cuticle can prevent the ants from escaping: during our retention trials, none of the ants managed to escape, and several were unable to pull themselves out of the fluid despite having a sufficient foothold on the glass wall (V.K., personal observations). Similarly, for a winged insect, any wetted body part will act like tethers to the fluid and further inhibit escape. Moreover, the reduced surface tension of PF causes insects to sink more readily, leading to larger areas of the cuticle being wetted and increasing the overall effort needed for escape. These two PF properties can act in synergy to facilitate prey retention.

### 4.3. Pitcher fluid resists dewetting and is difficult to remove

Perhaps the most striking but previously unrecognised PF behaviour was its strong resistance to dewetting from both hydrophilic and hydrophobic surfaces. When a droplet of PF was withdrawn from a glass surface, the initial contact line failed to move inward and instead the fluid formed a thin layer as it was actively withdrawn. PF also slowed down or prevented dewetting on hydrophobic surfaces, where it formed long fractal-like filaments branching out towards the initial rim. As a result, PF did not completely dewet on any of the tested surfaces, and large areas of the contact zone were left covered in PF. In contrast, water consistently dewetted on both hydrophilic and hydrophobic surfaces. Previous studies have shown that viscoelastic fluids can exhibit large wetting and dewetting contact angle hysteresis [63,64]. Moreover, a previous study demonstrated that shear-thinning viscoelastic fluids (aqueous solutions of high molecular weight polyacrylamide or polyethylene oxide) can produce films and filaments on hydrophobic surfaces and thereby resist dewetting over a greater range of retraction velocities than a Newtonian fluid (glycerin) [65]. We also observed filament deposition on hydrophobic surfaces with sticky *Nepenthes* PF, which is a shear-thinning viscoelastic fluid [13]. Such behaviour may enhance prey retention through the following mechanism: when an insect lands on the PF and struggles, parts of its body will become wetted. Through its struggles, the insect will raise its limbs or wings above the fluid surface (upstroke) then back down into the fluid (downstroke). According to our dewetting experiments, if the fluid was water, its limbs will dewet from the surface during the upstroke, and upon the downstroke, water will need to re-wet the surface, resulting in no overall advancement on the contact line. With PF, however, the upstroke will not dewet the fluid from the limb, and upon the downstroke, the fluid will readily interact with itself through the film or filament residues, thus facilitating the extension of the fluid contact line. In other words, the ant will be trapped in a positive feedback loop, akin to a ratchet motion, thereby constantly dragging itself into the fluid. Such a mechanism can work in combination with the reduced surface tension and may explain the sinking phenomena in PF but not in water. SEM images of tested gasters clearly show cuticular surfaces coated in PF residues, which is in stark contrast to the clean water-tested gasters. Analogous findings have been reported with carnivorous sundew plants (*Drosera* genus) that secrete viscoelastic glue-like mucilage from stalked glands to ensnare their prey. This mucilage readily spreads on lepidopteran wings and leaf surfaces that are highly non-wettable [66], and produces static contact angles lower than water on hydrophobic surfaces (47° compared to 83°) [67]. This implies that it is more energetically favourable for *Drosera* mucilage to interact with hydrophobic surfaces than water, which is analogous to our findings with *N. rafflesiana* PF. The delayed or prevented dewetting may be another important effect of the viscoelastic nature of the *N. rafflesiana* PF, which is likely a result of large (high molecular weight) acidic polysaccharides thought to be present within the fluid. Additional experiments are underway to characterise these polysaccharides and to examine the fractal-like filament deposition on hydrophobic substrates by a dilute biopolymer solution. Our findings highlight the potential for viscoelastic *Nepenthes* pitcher fluid to serve as a model for studying the mechanics of complex biological fluids.

## 5. Conclusions

Pitcher plants rely on multiple mechanisms to capture and retain insect prey. Aside from the well-studied structure-based mechanisms, the PF inside the trap serves both a digestive and a mechanical function for prey retention. We investigated how the sticky PF from *N. rafflesiana* adheres to insect cuticle and facilitates prey retention. Our findings reveal that PF surface tension is lower than water. This partly explains our observations of ants being wetted and sinking in the PF. Force measurements of insect cuticle dipped in and out of PF showed significantly greater work is required to retract from PF than from water. This arose from stable filaments between the cuticle and the PF that produced a force back towards the fluid. Image analysis of the tested cuticular surfaces confirmed that PF remained on the cuticle. Furthermore, PF resisted dewetting on insect cuticle as well as hydrophobic and hydrophilic surfaces, and left residues that could facilitate subsequent re-wetting of the cuticle. On the basis of our results, we propose that prey retention is based on a combination of three mechanisms: (1) when an insect falls into the pitcher and lands on the fluid, it readily breaks through the meniscus; (2) once wetted, the fluid’s resistance to dewetting prevents the insect from successfully separating itself from the liquid; (3) it takes the insect more effort to escape due to the formation of filaments that pull its body back into the fluid. Repeated attempts to escape only lead to further wetting of the cuticle, eventually ending with the prey being trapped by complete submersion or exhaustion.

## Acknowledgements

We would like to thank R. Mashoodh for her assistance with the statistical analysis, K. Ho for helping with the trials on ant swimming behaviour, and K.H. Muller and J.N. Skepper at the Cambridge Advanced Imaging Centre for their help in preparing and imaging SEM samples. We are grateful to A. Summers and his colleagues at the Cambridge University Botanical Gardens and U. Bauer at the University of Bristol for giving us access to their *Nepenthes* collections. We thank V. G. Baeza for providing the watercolour drawing of *N. rafflesiana* for the graphical abstract.

## Data availability

Data for the surface tension measurements are available in the Supplementary Materials. Data from the force measurements will be uploaded to Mendeley Data prior to publication.

