## Supplementary material for "How a sticky fluid facilitates prey retention in a carnivorous pitcher plant (*Nepenthes rafflesiana*)": supplemementary_materials

### Supplementary materials

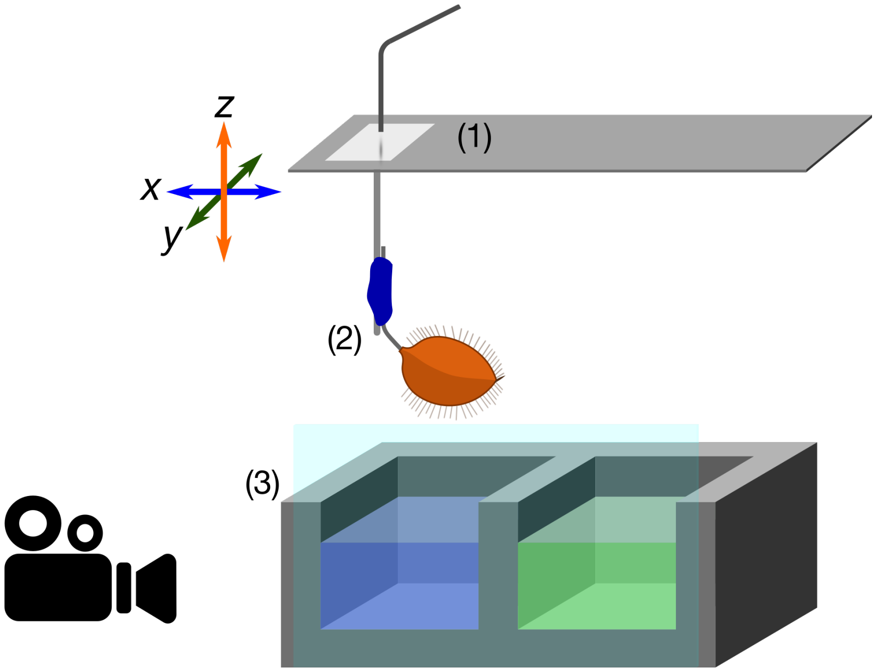


Supplementary Figure 1: Schematic of the fibre-optic force transducer set-used to quantify forces acting on ant gaster upon retraction from fluids. (1) A fibre optic sensor signal reflecting off the reflective metal foil glued on a thin metal beam was used as the translate displacement into forces. (2) An ant gaster mounted on an insect pin (diameter 0.1 mm) was attached to the force transducer using dental wax. The metal beam and the gaster could be moved in 3-dimensions with a motor stage. (3) A 3D printed aquarium with two contiguous chambers (each 13.7 x 9.1 x 8 mm in width x depth x height), one for water (blue) and the other for pitcher fluid (green). After testing in water, the gaster was moved to pitcher fluid and tested without needing to re-mount the specimen. Video recordings were synchronised with the motor stage movements.

**Additional details on calculating the work of retraction**

From each gaster trial, we calculated the 'work of retraction' using the following equation:

$$W_{retraction} = \sum_{i=a}^{b} F(z_i) \cdot \Delta z_i$$

where $F(z_i)$ is the force measured at a specific motor position $z_i$, $\Delta z_i$ is the distance moved between two force measurements, and *a* and *b* denote the start and end indices for the summation. *a* was defined as the index when the first positive force value was recorded. The second index, *b,* depended on the test liquid: for water, this was defined as when the gaster-liquid bridge suddenly collapsed (based on the synchronised video recordings). For PF, if the force-trace returned to zero before the next motor movement, the index immediately preceding this was used; if there was no negative force value by the end of the motor movement, *b* was the index of the last force value of the upward movement.

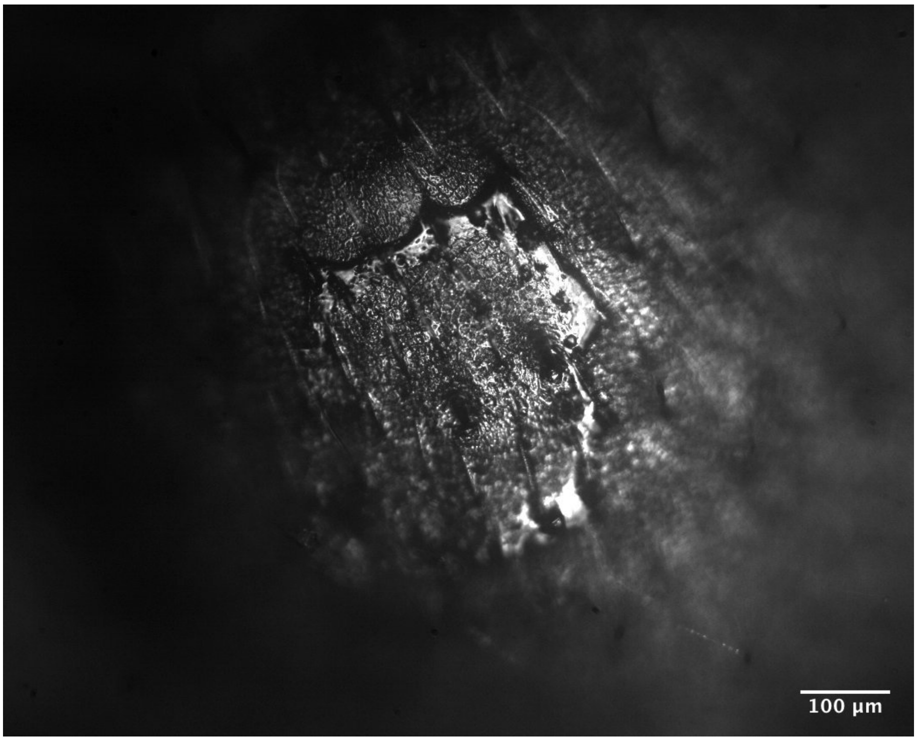


Supplementary Figure 2: Pitcher fluid does not easily dewet from inset cuticle, as seen in this light microscopy image of an ant gaster after a droplet of sticky pitcher fluid was placed on the surface and left to dry.

**Captions to supplementary videos**

Supplementary Video 1: Example videos demonstrating ant retention in *N. rafflesiana* pitcher fluid compared to water. Note: these trials were not the ant retention trials as described in the main text, and thus only two outcomes are shown ("walking on water" and "sunken").

Supplementary Video 2: Video-recording of a water droplet dewetting from a clean glass surface as observed under interference reflection microscopy.

Supplementary Video 3: Video-recording of *N. rafflesian* pitcher fluid dewetting from a clean glass surface as observed under interference reflection microscopy.

Supplementary Video 4: Video-recording of a water droplet dewetting and evaporating from the cuticular surface of an ant gaster (*Atta cephalotes*).

**Supplementary Table 1: Surface tension measurement values including Worthington numbers**

| SAMPLE | BOND NUMBER | WORTHINGTON NUMBER | SURFACE TENSION | VOLUME (µL) |
|---|---|---|---|---|
| **AP1** | 0.26 | 0.78 | 59.55 | 11.86 |
| **AP1** | 0.33 | 0.77 | 51.64 | 15.21 |
| **AP1** | 0.32 | 0.74 | 55.69 | 15.83 |

| | | | | |
|---|---|---|---|---|
| **AP1** | 0.33 | 0.73 | 47.74 | 13.36 |
| **AP1** | 0.25 | 0.72 | 60.26 | 11.14 |
| **AP2** | 0.33 | 0.78 | 50.54 | 15.06 |
| **AP2** | 0.33 | 0.77 | 53.13 | 15.64 |
| **AP2** | 0.32 | 0.77 | 55.70 | 16.37 |
| **AP2** | 0.32 | 0.76 | 53.72 | 15.71 |
| **AP2** | 0.32 | 0.72 | 52.77 | 14.59 |
| **AP3** | 0.31 | 0.77 | 65.05 | 19.19 |
| **AP3** | 0.32 | 0.76 | 59.36 | 17.30 |
| **AP3** | 0.31 | 0.76 | 65.84 | 19.17 |
| **AP3** | 0.32 | 0.76 | 56.10 | 16.32 |
| **AP3** | 0.31 | 0.76 | 65.41 | 18.97 |
| **AP4** | 0.32 | 0.77 | 56.80 | 16.80 |
| **AP4** | 0.33 | 0.77 | 51.52 | 15.24 |
| **AP4** | 0.32 | 0.77 | 56.23 | 16.63 |
| **AP4** | 0.32 | 0.77 | 55.07 | 16.26 |
| **AP4** | 0.32 | 0.77 | 60.60 | 17.87 |
| **AP4BRI** | 0.27 | 0.81 | 62.12 | 12.92 |
| **AP4BRI** | 0.26 | 0.81 | 66.70 | 13.85 |
| **AP4BRI** | 0.26 | 0.80 | 68.04 | 13.93 |
| **AP4BRI** | 0.26 | 0.80 | 66.20 | 13.54 |
| **AP4BRI** | 0.26 | 0.80 | 65.30 | 13.27 |
| **AP5** | 0.26 | 0.74 | 58.50 | 11.14 |
| **AP5** | 0.25 | 0.74 | 64.28 | 12.15 |
| **AP5** | 0.24 | 0.67 | 63.02 | 10.79 |
| **AP5** | 0.24 | 0.66 | 61.52 | 10.32 |
| **AP5** | 0.24 | 0.64 | 59.11 | 9.65 |
| **AP5-BRI** | 0.26 | 0.80 | 68.56 | 13.95 |
| **AP5-BRI** | 0.26 | 0.79 | 68.05 | 13.69 |
| **AP5-BRI** | 0.32 | 0.79 | 63.39 | 19.11 |
| **AP5-BRI** | 0.31 | 0.78 | 63.69 | 18.99 |
| **AP5-BRI** | 0.32 | 0.77 | 62.08 | 18.41 |
| **GP1-BRI** | 0.32 | 0.77 | 54.53 | 16.15 |
| **GP1-BRI** | 0.32 | 0.77 | 57.13 | 16.90 |
| **GP1-BRI** | 0.31 | 0.76 | 62.32 | 18.16 |
| **GP1-BRI** | 0.31 | 0.74 | 59.44 | 16.79 |
| **GP1-BRI** | 0.30 | 0.67 | 59.12 | 15.23 |
| **GP3-BRI** | 0.27 | 0.81 | 58.37 | 12.08 |
| **GP3-BRI** | 0.27 | 0.81 | 60.05 | 12.41 |
| **GP3-BRI** | 0.27 | 0.80 | 58.41 | 11.99 |
| **GP3-BRI** | 0.27 | 0.80 | 55.84 | 11.36 |
| **GP3-BRI** | 0.27 | 0.79 | 58.13 | 11.73 |
| **GP4-BRI** | 0.26 | 0.81 | 68.57 | 14.15 |
| **GP4-BRI** | 0.26 | 0.80 | 69.37 | 14.17 |
| **GP4-BRI** | 0.26 | 0.80 | 69.55 | 14.20 |
| **GP4-BRI** | 0.26 | 0.79 | 67.76 | 13.76 |
| **GP4-BRI** | 0.25 | 0.78 | 68.04 | 13.58 |
| **WATER** | 0.25 | 0.80 | 72.62 | 15.07 |
| **WATER** | 0.25 | 0.80 | 72.79 | 15.11 |
| **WATER** | 0.26 | 0.80 | 71.81 | 14.89 |
| **WATER** | 0.26 | 0.80 | 71.13 | 14.74 |
| **WATER** | 0.26 | 0.80 | 71.82 | 14.86 |
| **WATER** | 0.25 | 0.80 | 72.11 | 14.87 |
| **WATER** | 0.25 | 0.80 | 73.17 | 15.08 |
| **WATER** | 0.25 | 0.80 | 72.97 | 15.03 |
| **WATER** | 0.25 | 0.80 | 72.23 | 14.87 |
| **WATER** | 0.25 | 0.79 | 72.06 | 14.80 |

| | | | | |
|---|---|---|---|---|
| **XG 0.5%** | 0.35 | 0.84 | 53.41 | 14.25 |
| **XG 0.5%** | 0.35 | 0.81 | 52.84 | 13.65 |
| **XG 0.5%** | 0.36 | 0.89 | 53.78 | 15.20 |
| **XG 0.5%** | 0.35 | 0.83 | 53.45 | 14.07 |
| **XG 0.5%** | 0.35 | 0.85 | 54.62 | 14.78 |
| **XG 0.2%** | 0.32 | 0.73 | 60.36 | 15.76 |
| **XG 0.2%** | 0.32 | 0.74 | 59.84 | 15.79 |
| **XG 0.2%** | 0.32 | 0.75 | 60.51 | 16.12 |
| **XG 0.2%** | 0.32 | 0.73 | 59.21 | 15.52 |
| **XG 0.2%** | 0.32 | 0.74 | 60.29 | 15.92 |
| **XG 0.1%** | 0.31 | 0.75 | 66.78 | 18.48 |
| **XG 0.1%** | 0.31 | 0.75 | 66.59 | 18.53 |
| **XG 0.1%** | 0.30 | 0.72 | 66.73 | 17.68 |
| **XG 0.1%** | 0.31 | 0.73 | 66.77 | 18.04 |
| **XG 0.1%** | 0.31 | 0.74 | 66.89 | 18.36 |
